# Measuring the impact of genetic heterogeneity and chromosomal inversions on the efficacy of CRISPR-Cas9 gene drives in different strains of *Anopheles gambiae*

**DOI:** 10.1101/2023.03.31.535088

**Authors:** Poppy Pescod, Giulia Bevivino, Amalia Anthousi, Ruth Shelton, Molly Margiotta, Josephine Shepherd, Fabrizio Lombardo, Tony Nolan

**Affiliations:** Vector Biology Department, Liverpool School of Tropical Medicine, Liverpool, UK; Department of Public Health and Infectious Diseases, Division of Parasitology, University of Rome “la Sapienza”, Rome, Italy; Department of Biology, University of Crete, Vassilika Vouton, 71409 Heraklion, Crete, Greece; Institute of Molecular Biology and Biotechnology, Foundation for Research and Technology-Hellas, Heraklion, Greece; Department of Life Sciences, Imperial College London, UK

**Keywords:** Gene drive, malaria, *Anopheles*, genetic variation

## Abstract

The human malaria vector *Anopheles gambiae* is becoming increasingly resistant to insecticides, spurring the development of genetic control strategies. CRISPR-Cas9 gene drives can modify a population by creating double-stranded breaks at highly specific targets, triggering copying of the gene drive into the cut site (‘homing’), ensuring its inheritance. The DNA repair mechanism responsible requires homology between the donor and recipient chromosomes, presenting challenges for the invasion of lab-developed gene drives into wild populations of target species *An. gambiae* species complex, which show high levels of genome variation.

Two gene drives (*vas2*-5958 and *zpg*-7280) were introduced into three *An. gambiae* strains collected across Africa with 5.3-6.6% variation around the target sites, and the effect of this variation on homing was measured. Gene drive homing across different karyotypes of the 2La chromosomal inversion was also assessed. No decrease in gene drive homing was seen despite target site heterology, demonstrating the applicability of gene drives to wild populations.

## Introduction

### Gene drives in vector control

Global control efforts have averted an estimated 1.5 billion cases of malaria in the last two decades but this progress has begun to slow, with 619 000 deaths reported in 2021 alone ^1^. Malaria transmission persistence has been attributed to a combination of stalling or inadequate control programs, insecticide resistance of the mosquito vector, and treatment resistance of the parasite ^2^. The World Health Organisation has stressed the importance of developing novel control strategies to meet its malaria elimination goals ^3,4^.

Genetic control strategies can achieve population modification or suppression of a target species without collateral damage to non-targets or the environment by the modification of the target genome, making these strategies highly desirable alternatives to widespread insecticide use. One such strategy is the use of selfish genetic elements with super-Mendelian inheritance rates known as gene drives. Gene drives can deliver a genetic payload or disrupt an essential gene while overcoming any subsequent fitness cost by severely biasing its own inheritance, allowing its spread in a population ^5-10^. Strategies using gene drives are being considered for the control of several pest species ^11,12^, and have progressed to the successful development of CRISPR/Cas9-based gene drives in the primary vector of malaria in Africa, *Anopheles gambiae* ^7^.

The Cas9 protein guided by an sgRNA is capable of making highly-specific double-stranded breaks (DSBs) in a chromosome, allowing the introduction of an alternate sequence at the cut site using the cell’s own DNA repair mechanism ^7^. DSB repair by the cell can involve the use of a homologous template strand, usually the paired chromosome, which is copied to accurately repair the break ^13,14^. When a gene drive element is copied into the broken chromosome alongside the homologous template sequence, the gene drive is inserted at the breakpoint in a process known as homing.

Homing from one chromosome to another in germline cells means the gene drive will be integrated in the majority of gametes, resulting in its super-Mendelian inheritance in the next generation. This mechanism can be exploited to bias the inheritance of a coupled effector gene through a population, such as antimicrobial peptides to impede malaria development ^10^, or to target a gene essential for fertility and therefore reduce the target population size ^7^.

### Gene drive resistance

The emergence of resistance to gene drives has been demonstrated empirically in synthetic drive constructs ^15,16^. CRISPR-based gene drive resistance occurs as small genetic differences at the cut site, reducing gRNA binding and therefore the ability of the Cas9 enzyme to create a DSB. These cut site mutations can arise during the DSB repair process via alternate repair pathways such as non-homologous end-joining, which enzymatically repairs the cut without a template but with higher rates of error ^15,17-19^. If these genetic differences produce a functional allele with a fitness cost less than that of the gene drive, they may be positively selected for in the population. Functional alternate alleles produced by gene drive-induced mutations can reduce CRISPR-gRNA binding enough to confer complete resistance to the gene drive ^15^.

Strategies to reduce the likelihood of resistance developing have been suggested; modern gene drives will target genes which are haplosufficient (one functional copy is required for survival or fertility) and highly conserved, therefore making any mutations at the target site likely to result in unviability ^8,15,20^. This reduces the speed of gene drive resistance development but does not entirely prevent it; mutations produced during non-homologous end joining (NHEJ) will still eventually lead to resistance ^15^. NHEJ, and therefore related mutations conferring resistance, can be reduced by using more efficient germline-specific promoters with less accidental somatic expression of the Cas9 enzyme ^21^. Multiple target sites in different genes can be used in a single gene drive system by multiplexing gRNAs; homing can occur at all target sites, making independently-evolved resistance at all target sites necessary to prevent super-Mendelian inheritance of the gene drive ^22-24^.

Any intervention which applies a strong selection pressure will eventually produce a similarly strong and concomitant pressure to evolve resistance. Resistance has historically only been discovered after the implementation of a control strategy, and after the resistance has become a public health issue ^25^. By anticipating and investigating potential issues such as resistance during gene drive development we can reduce the impact on control strategies. Genetic variation at the target site, whether produced by Cas9-mediated NHEJ or naturally present in a target population, could act as a barrier to successful implementation of gene drives.

### Genetic variation – a barrier to gene drive success?

#### Single nucleotide polymorphisms

Genetic differences around a gene drive target site, or target locus heterology (TLH), may occur naturally in a wild population even in highly functionally constrained genes. Differences within the gRNA target site have the most impact on drive efficiency ^26^, but due to the nature of DSB repair TLH will also potentially reduce the gene drive homing rate. Stringent regions of homology between the donor chromosome (containing the region to be copied) and the recipient chromosome (where the DSB occurs) are required for homology-directed repair (HDR) in mammalian cells, where 1.2% TLH within 1kb of the DSB causes a 22% reduction in the recombination required for HDR ^27^. Similar dependence on homology has been noted in *Drosophila melanogaster*, where 1.4% TLH suppressed recombination by 32% ^28^; and *Aedes aegypti*, with 1.2% TLH created by experimental recoding resulting in a 66% reduction in homing ^29^. Given the conserved nature of DNA repair mechanisms, it is reasonable to expect that this sequence homology requirement would extend to *An. gambiae*, which has an incredibly diverse genome including more than 57 million single-nucleotide polymorphisms (SNPs) ^30^.

#### Chromosomal inversions

In addition to SNPs, the *An. gambiae* species complex contains over 120 chromosomal inversions ^31,32^. These inversions vary in their geographical and seasonal distribution and have been associated with desiccation resistance, larval habitat, insecticide resistance, and malaria infection rate ^31,33-37^. The largest inversion in *An. gambiae* is the 2La/2L+^a^, which spans roughly half the length of chromosome 2L ^38^; the ancestral 2La form is implicated in anthropophilic behaviour, aridity tolerance, and *Plasmodium* transmission ^35,39-41^. Recombination of inverted chromosomes in opposite orientations is reduced as the chromosomes are forced to form a loop in order to align ^42^. Reduced recombination between inversion heterokaryotypes has been empirically demonstrated during meiosis in multiple species, including *Drosophila* (7.7-fold decrease) ^43^. The effect extends beyond the inversion breakpoints to suppress recombination in regions close to the inversion and increase recombination at distant regions, known as the interchromosomal effect ^44^, and can change the recombination landscape enough to suppress recombination in homokaryotypes as well ^45,46^.

A reduced recombination rate between inversion heterokaryotypes could theoretically interfere with HDR in gene drive releases, leading to reduced spread of a gene drive situated within the inversion into heterogenous wild populations. In an allelic drive system in *Drosophila*, inversion heterokaryotypes had a drive rate a third lower than inversion homokaryotypes ^47^. Meiotic recombination between 2La/2L+^a^ heterokaryotypes is at least 5-fold less than between 2L+^a^ homokaryotypes ^48^. However, multiple gene drive systems have been developed within the 2La inversion site in *An. gambiae* successfully, with super-Mendelian inheritance (76-98%) ^49,50^. As these were not developed with the 2La inversion karyotype in mind, or tested with different karyotypes, the impact on recombination rate during gene drive homing in *An. gambiae* is still unknown.

#### Variation outside of the target region

Genetic variation outside of the target region can also influence gene drive inheritance and resistance development. In a study of gene drive homing and resistance rates in different strains of *Drosophila*, all with identical target site sequences, inheritance rates ranged from 64.1-85.9%; increased inheritance was significantly associated with certain genotypes, but no SNPs were identified as contributing significantly ^51^. Moderate variation has been noted in gene drive homing in different genetic backgrounds of *Drosophila* despite little to no variation within 200 bp of the cut site ^19^. Background genetic variation can also influence the development of resistant alleles at the target site; no single gene was found to be significantly responsible for increased resistance development, indicating a combined effect of multiple genes ^51^. Differences in homing efficiency may be due to differences in a combination of genes, such as DNA repair mechanisms, DNA transcription or translation, or germline expression. In naturally occurring gene drives, suppressors can evolve to reduce the impact of the drive in the population; these are often unlinked to the gene drive, such as small RNAs or alterations in heterochromatin structure ^52^. Undoubtedly, the interaction between genetic variation and gene drive homing needs to be explored for their effective use in control strategies.

To assess the impact of TLH and inverted chromosomes on the homing of a gene drive element in *An. gambiae*, two well characterised lab-created gene drive strains *vas2*-5958 and *zpg*-7280 were crossed with three alternate *An. gambiae* wild type strains from across East, Central and West Africa (Kisumu, N’Gousso and Tiassale), all with TLH around the cut sites. The *vas2*-5958 gene drive element is located within the 2La inversion; gene drive homing rates were compared between 2La heterokaryotypes and homokaryotypes.

## Materials and Methods

### Mosquito rearing

All mosquitoes were reared under standard conditions of 26 ± 2°C and 70 ± 10% relative humidity, with a 12 hour light/dark cycle with one hour dusks/dawns. Larvae were fed on ground fish food flakes (TetraMin® tropical flakes) and adults were fed 10% sucrose solution *ad libitum*. Adults were allowed to mate for 5-10 days before blood feeding and egg collection.

### Mosquito strains

Two G3 colonies containing gene drive elements were used, both created by Hammond *et al*. and well characterised ^7,21^. The *vas2*-5958 colony contains a CRISPR/Cas9 endonuclease construct within AGAP005958, an ortholog of the *Drosophila yellow-g* gene expressed in somatic ovarian follicle cells with an unknown function ^53^. The AGAP005958 gene is located within the 2La inversion ^54^, with a gRNA cut site within 4Mb of the distal breakpoint. The *zpg*-7280 colony contains a similar construct in AGAP007280, ortholog of the *Drosophila nudel* gene also expressed in follicle cells with a known role in dorsoventral patterning of the developing embryo ^55^. Both genes have a haplosufficient role in female fertility, making them useful targets for population modification or suppression gene drive strategies.

The inserted construct for both colonies consists of: a CRISPR/Cas9 protein under germline-only promotion (*zpg* in the *zpg*-7280 line and *vas2* in the *vas2*-5958 line), a gRNA sequence targeting the cut site for each line respectively under U6 (universal) promotion, and a red fluorescence protein marker with a 3xP3 promoter, all flanked by *attB* recombination sites to allow insertion into previously created docking lines via recombinase-mediated cassette exchange ^56^. The full sequence of vector p165 used to produce these two lines, with the only difference between them the gRNA sequence, is available on GenBank (accession ID: KU189142) ^7^.

The wild type strains used in crosses were taken from colonies kept at the Liverpool School of Tropical Medicine ^7,57-59^; details can be found in Table 1.

**Table 1.** Wild type *Anopheles* strains used in this work.

| Strain | Species | Place and date of collection | Reference |
| --- | --- | --- | --- |
| G3 | *An. gambiae*/*An. coluzzii* hybrid | The Gambia, 1975 | (7) |
| N’Gousso | *An. coluzzii* | Yaoundé, Cameroon 2002 | (57) |
| Tiassale | *An. gambiae*/*An. coluzzii* hybrid | Cote d’Ivoire, 2012 | (58) |
| Kisumu | *An. gambiae* | Kenya, 1975 | (59) |

### Crosses of gene drive strains into alternate backgrounds

An outline of the methodology can be seen in Figure 1. The number and sex of mosquitoes used in each cross can be seen in Table 2. All F_1_ hybrid adults used in crosses were confirmed to be heterozygous for the gene drive element by screening for the RFP marker via fluorescent microscopy during the larval stage. Females containing the *vas2*-5958 gene drive are sterile due to unintended somatic promotion of Cas9 under the *vas2* promoter ^7^; therefore, in *vas2*-5958 crosses only males containing the gene drive were used. For *zpg*-7280 F_1_ crosses female hybrids were used. F_1_ cross females were forced to lay in single deposition and up to 50 offspring per female were screened for the presence of the RFP marker to determine the rate of gene drive in the hybrid parent. Drive rates were compared to data from Hammond *et al*. ^7,21^ using a pairwise Wilcoxon test with false discovery rate correction (Table S1).

**Table 2.**
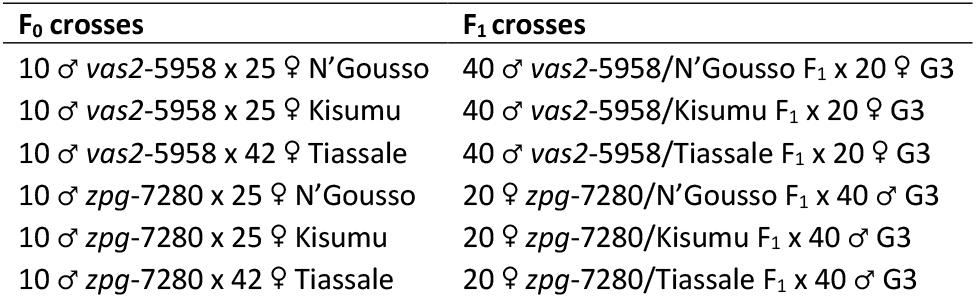
Details on the number, sex and strain of each F0 and F1 cross.

| F<sub>0</sub> crosses | F<sub>1</sub> crosses |
| --- | --- |
| 10 ♂ *vas2*-5958 x 25 ♀ N’Gousso | 40 ♂ *vas2*-5958/N’Gousso F<sub>1</sub> x 20 ♀ G3 |
| 10 ♂ *vas2*-5958 x 25 ♀ Kisumu | 40 ♂ *vas2*-5958/Kisumu F<sub>1</sub> x 20 ♀ G3 |
| 10 ♂ *vas2*-5958 x 42 ♀ Tiassale | 40 ♂ *vas2*-5958/Tiassale F<sub>1</sub> x 20 ♀ G3 |
| 10 ♂ *zpg*-7280 x 25 ♀ N’Gousso | 20 ♀ *zpg*-7280/N’Gousso F<sub>1</sub> x 40 ♂ G3 |
| 10 ♂ *zpg*-7280 x 25 ♀ Kisumu | 20 ♀ *zpg*-7280/Kisumu F<sub>1</sub> x 40 ♂ G3 |
| 10 ♂ *zpg*-7280 x 42 ♀ Tiassale | 20 ♀ *zpg*-7280/Tiassale F<sub>1</sub> x 40 ♂ G3 |

**Figure 1.**
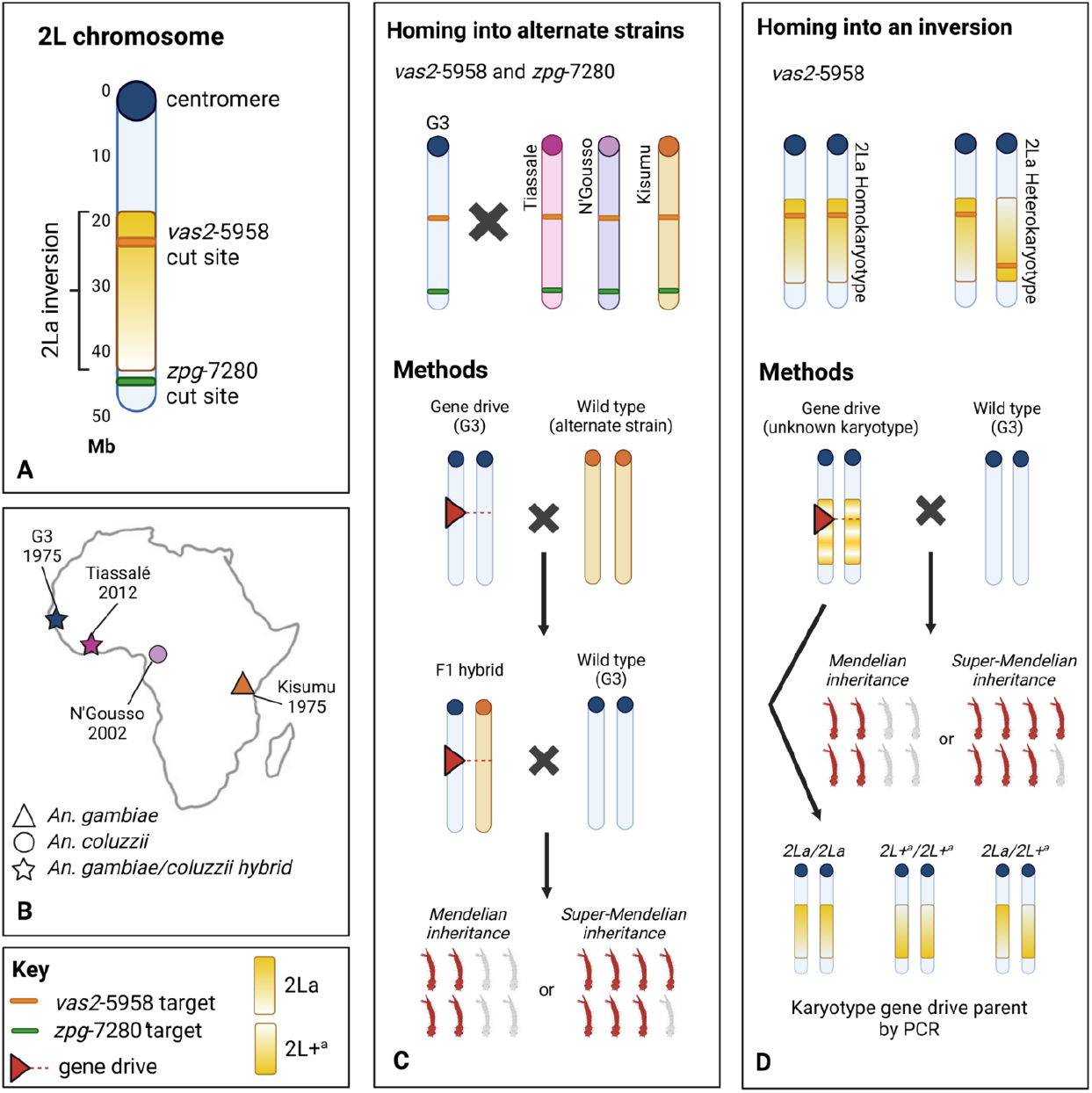
Experimental homing of a gene drive into alternate genetic backgrounds. A. The location of gene drive cassettes in the 2L chromosome of gene drive strains *vas2*-5958 and *zpg*-7280, relative to the centromere and the 2La inversion region, which is shown in wild type orientation. B. The locations and dates of capture for each strain, and their species identification. C. Methods for assessing gene drive homing into alternate strains. Gene drives were crossed with alternate strains to produce hybrid F_1_ individuals; these were backcrossed to G3 and the F_2_ offspring were screened for the RFP-tagged gene drive via fluorescent microscopy to determine the homing rate in the F_1_ hybrid. D. Methods for assessing gene drive homing in different inversion karyotypes. *vas2*-5958 gene drive individuals were crossed to wild type G3, the gene drive parent’s 2La inversion karyotype was determined by PCR, and the offspring screened for inheritance of the gene drive to investigate the impact of chromosomal inversion on gene drive homing rate.

### Target site sequence heterology

To determine the maximum potential TLH in each strain, F1 hybrids of each type were pooled and their wild type chromosome (representing each wild type strain) was sequenced. DNA was extracted from pools of 33-37 adults using a Wizard® genomic DNA purification kit (Promega) and a region of ∼690bp spanning the gRNA cut sites for both *vas2*-5958 and *zpg*-7280 gene drives was amplified in two fragments either side of the gene drive insert. Fragments were amplified by PCR using Phusion Hot Start II High-Fidelity DNA polymerase (Thermo Scientific™), with forward and reverse primers at a final concentration of 0.5 μM (Table S2) and 1 μl genomic DNA in a 50 μl reaction. PCR conditions were: an initial denaturation step at 98°C for 30 seconds; followed by 30 cycles of denaturation at 98°C for 30 seconds, 30 seconds at the annealing temperature (Table S2), and extension at 72°C for 15 seconds; and a final extension step of 10 minutes at 72°C.

PCR products were sequenced by Illumina MiSeq sequencing; reads were quality filtered and aligned against an amplicon containing all SNP variants present in G3 deep sequencing data ^15^ using CRISPResso ^60^. Alleles present at >1% relative abundance were aligned to G3 sequences in Benchling to determine the percentage of mismatch between G3 and each strain at the homing sites (raw files accession: PRJNA914102). As *vas2*-5958 is known to produce ‘leaky’ promotion and therefore maternal deposition of the Cas9 enzyme, resulting in NHEJ-induced deletions at the cut site in somatic tissue, any characteristic NHEJ deletions around the cut site in these F1 hybrids were removed from the TLH analysis.

### Homing rate analysis in alternative 2La karyotypes

Mosquitoes from the *vas2*-5958 colony were backcrossed to G3 and offspring were screened for the gene drive marker; 65 F1 males were mated individually to three G3 females, with eggs collected from each group and screened for the gene drive element. Each male parent was karyotyped for the 2La inversion as previously described ^38^, and drive rates in heterozygotes and homozygotes of both karyotypes were compared using a Wilcoxon test.

## Results and discussion

### *An. gambiae* gene drives are robust to TLH

The *vas2*-5958 and *zpg*-7280 *An. gambiae* gene drive lines, originally made in the G3 background and targeting haplosufficient female fertility genes, were crossed into three different strains to create F_1_ hybrids which were backcrossed to wild type G3 to assess the F_1_ hybrid homing rate (see Figure 1c). The TLH around the cut sites was 5.3-6.6% between each strain and the gene drive background strain (G3), with significant variation between the left and right sides of both gene drive cut sites (Figure 2 and Figure 3). No SNPs were observed within the gRNA sequence; however, a SNP was commonly observed in the *N* base of the *zpg*-7280 -*NGG* PAM site (Figure 3). All F_1_ hybrids for both gene drive colonies produced super-Mendelian inheritance rates of the gene drive element (*vas2*-5958: 81.8-100%; *zpg*-7280: 92.0-100%), with no significant difference from the control (Figure 4). No reduction in larval production was observed in *zpg*-7280 hybrids (Figure S2), suggesting no loss of fertility.

**Figure 2.**
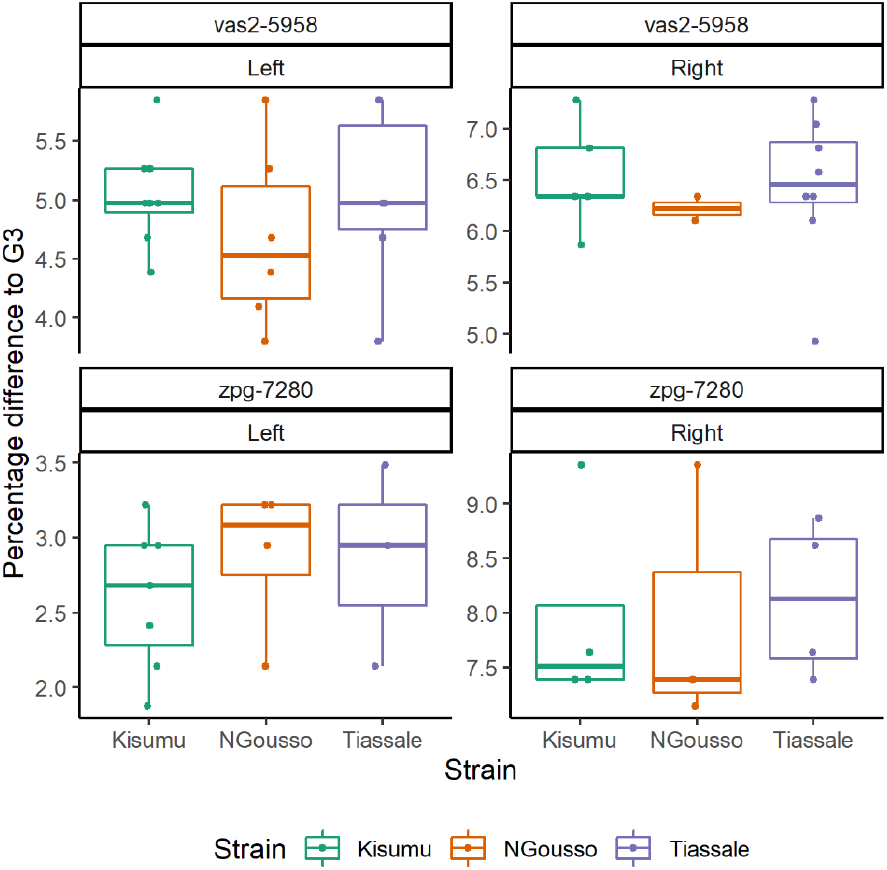
Target locus heterology in three strains (Kisumu, N’Gousso and Tiassale) compared to G3, at two gene drive sites (vas2-5958 and zpg-7280), with alleles present at >1% relative abundance. The data represents the maximum potential TLH between each strain and G3, by comparing each allele from the pooled F1 hybrid wild type chromosomes to a G3 sequence containing all known SNPS found in a deep sequencing dataset of 24 G3 individuals. Each point represents an allele from pooled sequencing of adult mosquitoes, with percentage difference to G3.

**Figure 3.**
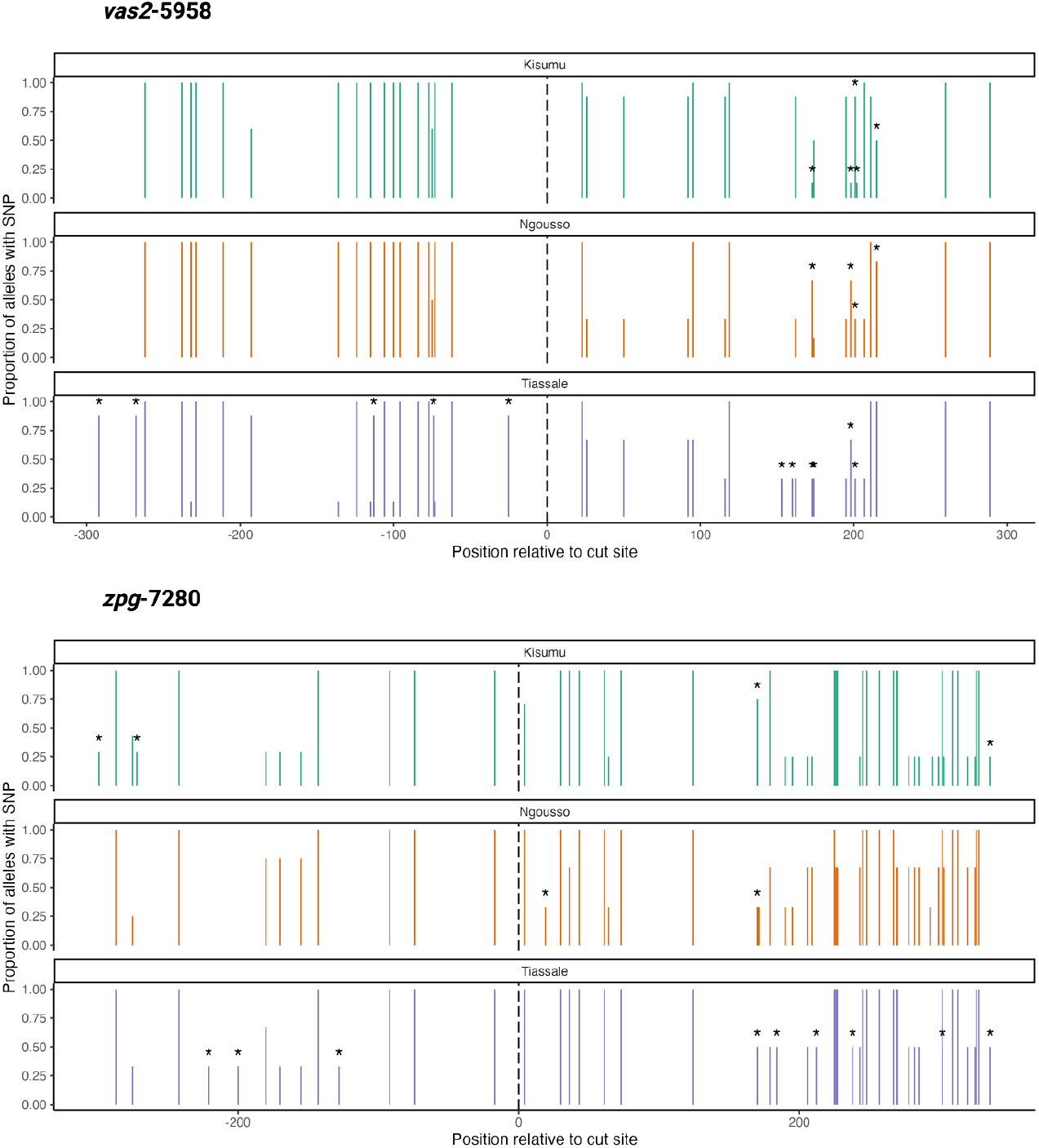
Position and frequency of SNPs at two gene drive sites (*vas2*-5958 and *zpg*-7280) in three strains (Kisumu, N’Gousso and Tiassale) compared to G3. The position of each SNP is given relative to the gene drive cut site, indicated by a dashed line. SNPs which are not also found in G3 are marked with an asterix. No SNPs were observed within the gRNA sequence of either cut site – however, at the *zpg*-7280 cut site a SNP commonly occurred in the *N* of the -*NGG* PAM site.

**Figure 4.**
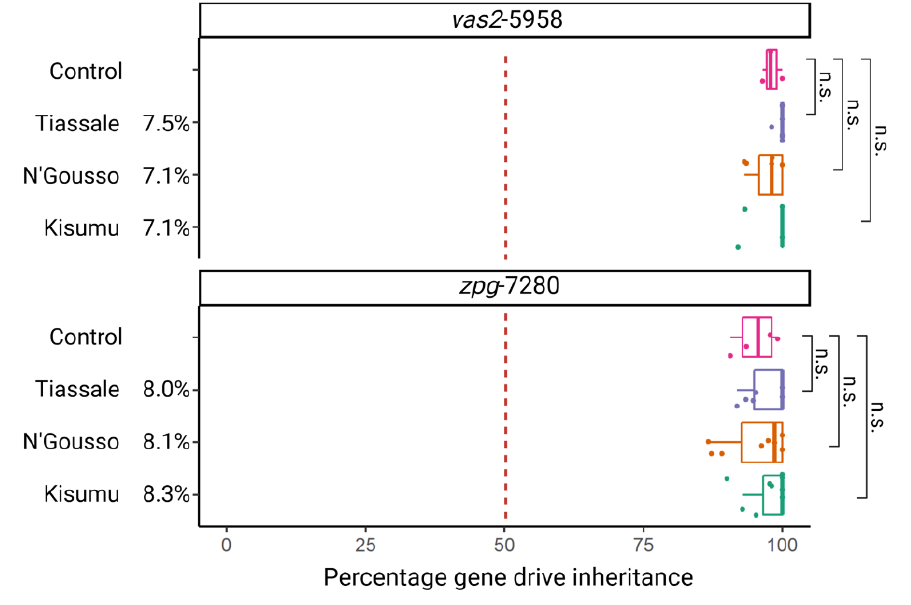
The inheritance rate of two gene drive elements vas2-5958 and zpg-7280 in the offspring of F1 hybrids of three different strains, compared to the control rate of inheritance in the gene drive colony (G3 background). Target locus heterology ∼600bp around the cut site between each strain and the G3 wild type is given in percentages next to strain names. Homing into alternate chromosomes produced drive rates which were not significantly different to the control drive rate (Pairwise Wilcoxon test, corrected for false discovery rate). n.s – non-significant.

This result varies considerably from previous findings in *Ae. aegypti* ^29^ and *D. melanogaster* ^28^, which saw significantly reduced HDR between sequences with 1.2% and 1.4% TLH respectively. Differences in methodology between the two studies and this work make direct comparison difficult; both used artificially generated silent mutations spaced at regular intervals to generate TLH, which could have a different impact recombination than the naturally occurring, irregularly-spaced TLH in the strains used here. Additionally, Ang *et al*. measured HDR between a donor plasmid and recipient chromosome rather than between chromosomes, and Do *et al*. used heat-inducible *I-Sce1* for DSB formation rather than Cas9. It may be the case that these previous studies, while well suited to describe their respective systems, were not good predictors of the dynamics of Cas9-based gene drive homing. While we cannot definitively state based on comparison to these studies that *An. gambiae* HDR is inherently more robust to TLH than *Ae. aegypti* or *D. melanogaster*, it appears that Cas9-based gene drive homing is efficient enough in *An. gambiae* that increased TLH is tolerated without causing enough of a reduction in efficiency to reduce the homing rate. This is supported by previous studies which have found Cas9-based gene drive homing rates are higher in *Anopheles* (∼97%)^8^ than in both *Drosophila* (∼80%)^19^ and *Aedes* (∼70%)^61^.

The robustness of *An. gambiae* gene drive homing to variation has important consequences for its application in real-world vector control strategies. The development of gene drives in lab-bred mosquito strains allows for standardisation of the genetic background for easier study but has called into question their applicability to heterogenous wild populations. Despite the sensitivity of homing in other organisms to low amounts of TLH, our findings show no significant reduction in homing activity into multiple strains with up to 6.6% TLH in *An. gambiae*. The strains used in this experiment were collected from East, Central and West Africa across a span of 37 years, and are a mixture of *An. gambiae, An. coluzzi* and *An. gambiae/An. coluzzi* hybrids (Figure 1b). The demonstration of unimpeded gene drive homing into strains of this diversity represents the strong potential for gene drive implementation across members of the *An. gambiae* species complex that are able to produce fertile progeny.

### No impact of 2La karyotype on gene drive homing

The *vas2*-5958 gene drive is located within the region covered by the 2La inversion (Figure 1a); homing rates for all three permutations of the 2La inversion were analysed (Figure 1d). There was no significant difference in homing rate between 2La inversion karyotypes (Figure 5).

**Figure 5.**
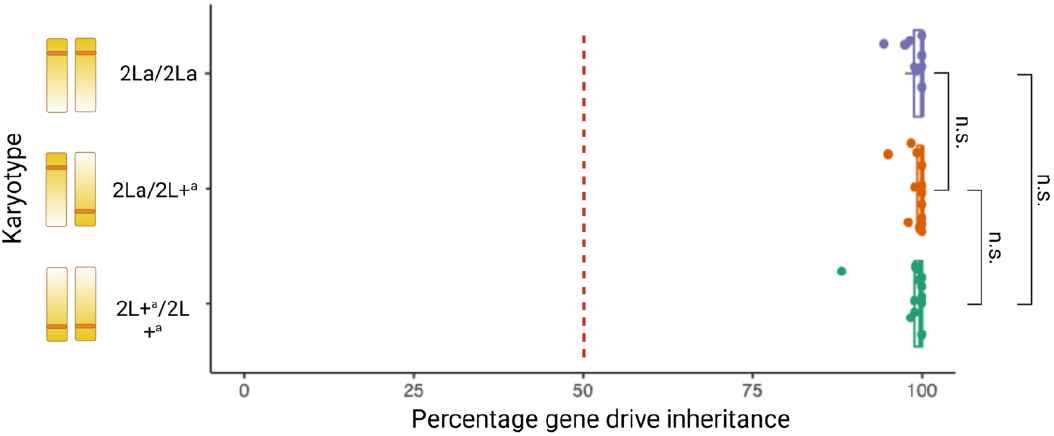
The inheritance rate of the *vas2*-5958 gene drive element in the offspring of males either homozygous (2La/2La and 2L+^a^/2L+^a^) or heterozygous (2La/2L+a) for the 2La chromosomal inversion. There was no significant difference in gene drive inheritance between the three karyotypes (Wilcoxon test). n.s. – non-significant.

Despite previous observations of reduced gene drive conversion across inversions in *Drosophila* ^47^ and reduced meiotic recombination within the 2La inversion region in *An. gambiae* ^48^ we saw no evidence of reduced gene drive homing rate in 2La inversion heterokaryotypes. However, reduced recombination is not uniform across an inversion, and adjacent sequences external to the inversion can also show altered recombination rates. Meiotic recombination is slightly higher in the middle of the inversion compared to regions near the breakpoints, due to the increased ease of forming chiasmata between sister chromatids at the centre of the inversion loop ^47,48^. The *vas2*-5958 gene drive target site is <4 Mb from the distal breakpoint of the 2La inversion (Figure 1a) ^7,62^; theoretically, recombination at this point should have been low, but this is not reflected in the gene drive rates we observed.

Adjacent to the inversion, the region between the proximal breakpoint of the 2La inversion and the centromere shows strong recombination reduction, with a less strong but still reduced recombination rate in the region distal to the centromere ^48^. The *zpg*-7280 gene drive target site is 2.8 Mb from the 2La distal breakpoint and is therefore located in a region with a known slight reduction in meiotic recombination ^7,62^. Our results suggest that this is not sufficient to reduce homing, but future work could explore if other targets within the inversion, or closer to the breakpoint, may be affected.

While homing does not appear to be reduced within the inversion in *An. gambiae*, other impacts of the inversion on long-term control strategies should be considered. Reduced meiotic recombination results in protection of the inverted regions and their accumulation within populations; a common mechanism of speciation in *Anopheles* ^54^. Linked regions can result in persistence of deleterious mutations or the spread of adaptive alleles for certain environments. In the case of gene drives, regardless of the impact of recombination on the homing mechanism itself, inversions could impact the penetrance of gene drives into wild populations indirectly, through reproductive isolation. That said, unless this reproductive isolation is total, even rare cases of intra-strain hybridisation should lead to the gene drive rapidly introgressing into the new karyotype. There is good precedent for this in the adaptive introgression of insecticide resistance alleles between *An. gambiae* and *An. coluzzii*, two separate species that are not fully isolated reproductively ^63^. The idea of ‘forced’ introgression, whereby gene drives are backcrossed into wild populations before release, has been suggested to reduce the introduction of novel chromosomal arrangements or variation into wild populations ^64^.

### Application of gene drives to wild populations

At first glance, the high level of variation in the *An. gambiae* species complex suggests that gene drives developed in lab-bred colonies could struggle to spread in wild populations via HDR. Our results suggest that this is not the case; with a highly conserved gRNA, variation in the surrounding sequence or in the chromosomal structure had no impact on the gene drive constructs tested here. The use of highly conserved gRNA sites is an important strategy for reducing the development of gene drive resistance ^8^. The availability of deep sequencing data for *An. gambiae* via the Ag1000G confirms the high variation within the species complex, but also greatly improves our ability to choose gRNAs appropriately ^65^. Correspondingly, gRNA target sites must be chosen carefully to confine gene drives to a particular strain; there are a variety of self-limiting strategies currently in development that either combine non-autonomous elements or target alleles private to the target population ^66-68^.

The specificity of the gRNA targeting system produces very low off target effects in *An. gambiae*, making CRISPR/Cas9 gene drives resistant to unexpected homing outside of the target sequence ^69^. However, there is potential for neighbouring sequences flanking the gene drive to be carried over during HDR due to resection of the broken chromosome ^28^. This could result in tight allelic linkage of neighbouring sequences to the gene drive and introgression of novel alleles into wild strains, suggesting that gRNA target regions need to be chosen with the surrounding sequences in mind.

Future work will be able to determine the precise dynamics of genetic exchange between the gene drive donor and recipient chromosome.

Regardless of TLH of up to 6.6%, gene drive strategies for *An. gambiae* control show promising efficacy for malaria control in wild mosquito populations. The self-sufficiency of gene drives after initial release has meant extra care is being taken to characterise how gene drives will function in natural settings ^70,71^. This work offers improved understanding of gene drive dynamics in wild populations and demonstrates their potential for *Anopheles* control.

## Authorship contribution statement

**Poppy Pescod**: Methodology, Software, Validation, Formal analysis, Investigation, Data curation, Writing – Original Draft, Writing – Review & Editing, Visualization. **Giulia Bevivino**: Methodology, Investigation, Writing – Review & Editing. **Amalia Anthousi**: Methodology, Validation, Investigation, Supervision, Writing – Review & Editing. **Ruth Shelton**: Investigation, Writing – Review & Editing. **Molly Margiotta**: Investigation, Writing – Review & Editing. **Josephine Shepherd**: Investigation, Writing – Review & Editing. **Fabrizio Lombardo**: Conceptualization, Methodology, Supervision, Project administration, Funding acquisition. **Tony Nolan**: Conceptualization, Methodology, Validation, Resources, Writing – Review & Editing, Supervision, Project administration, Funding acquisition.

## Supporting information

### S1 Appendix: Analysis of a fourth strain, Busia

In a separate piece of work, a fourth strain Busia (*Anopheles gambiae* s.s., captured in Uganda in 2018) (Lynd, *et al*., 2019) was analysed for homing rates in F1 hybrids with both *vas2*-5958 and *zpg-* 7280. Male hybrids of Busia with both gene drives were used in *en masse* crosses with wild type G3 females; females were put in single deposition for drive rate analysis (four from crosses with Busia/*vas2*-5958 hybrids, and 11 from crosses with Busia/*zpg*-7280 hybrids). Target site heterology (TLH) was calculated from wild type Busia using Sanger sequencing, with nine replicates on the left-hand side of the *vas2*-5958 cut site, and three replicates spanning 700 bp either side of the *zpg*-7280 cut site (accession: PRJNA914102). While the Busia samples were processed differently to the remaining three strains and were therefore left out of the main analysis, they show the same pattern of uninterrupted homing regardless of TLH, indicating the robustness of this effect to different analysis methods. All supporting information will include results from the Busia strain alongside Kisumu, N’Gousso and Tiassale.

Lynd *et al*. (2019). LLIN Evaluation in Uganda Project (LLINEUP): a cross-sectional survey of species diversity and insecticide resistance in 48 districts of Uganda. *Parasites & Vectors* **12**(1), 1-10.

**Figure S1:**
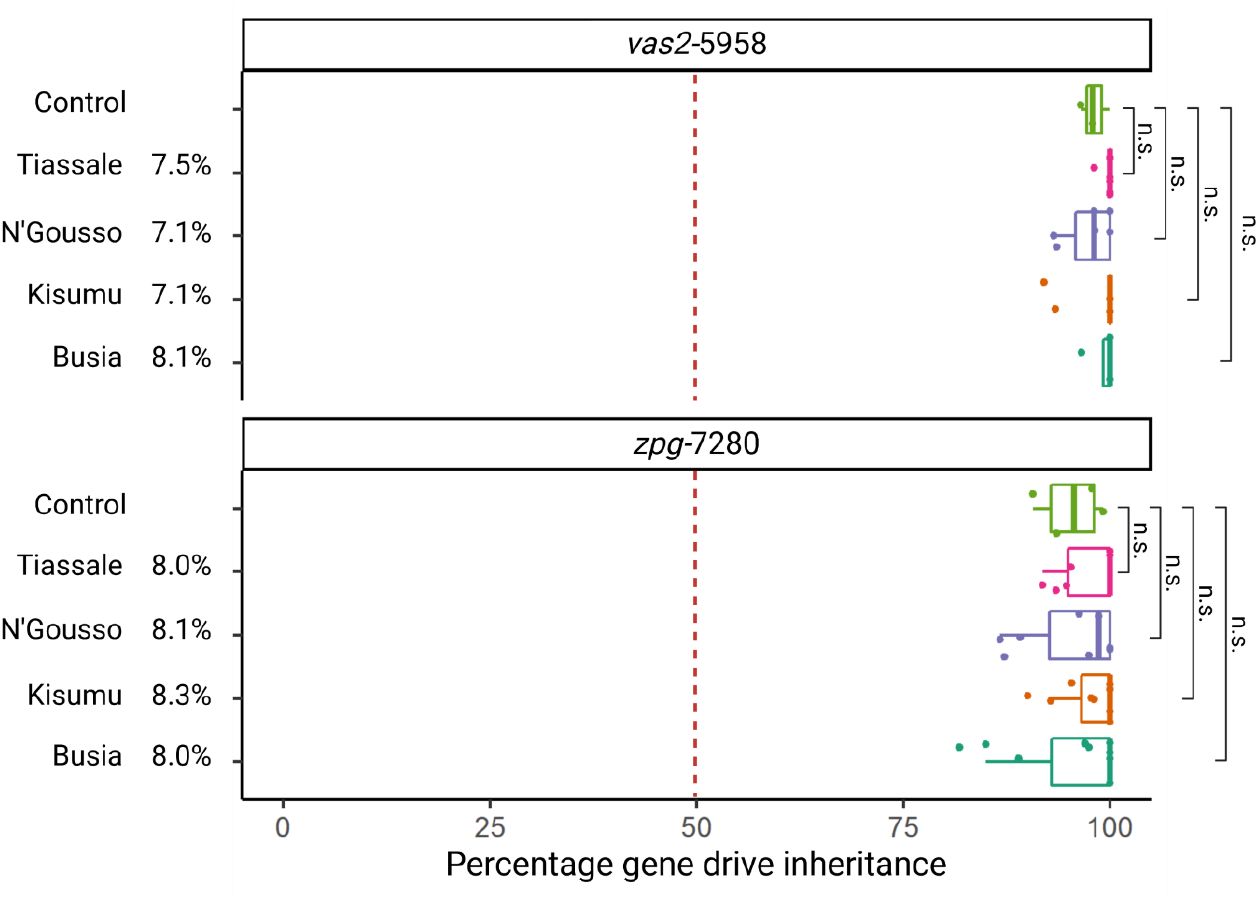
Gene drive inheritance rate analysis of Kisumu, N’Gousso, Tiassale and Busia hybrids with *zpg*-7280 and *vas2*-5958 gene drives. TLH is given as a percentage next to each strain name; statistical analysis was conducted using a pairwise Wilcoxon test with false discovery rate correction. n.s. – non-significant.

**Table S1:**
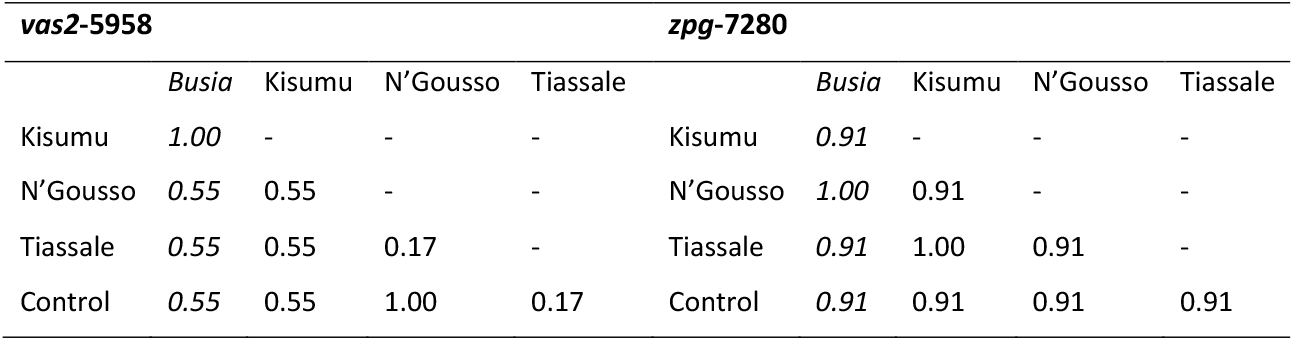
Pairwise Wilcoxon test p values with false discovery rate correction for gene drive inheritance rate. Comparisons were performed for hybrids with all four strains of both gene drives. No comparisons were statistically significant.

### S2 Appendix: Analysis of TLH between chromosomes within single *zpg*-7280/N’Gousso F1 hybrids

To corroborate our estimates of TLH from pools, a region spanning ∼700 bp either side of the cut site of six individual *zpg*-7280/N’Gousso F1 hybrids was sequenced for both chromosomes, allowing the calculation of exact TLH within each individual hybrid. DNA extractions from individual F1 adults and PCR reactions were carried out as in the main methodology; PCR products were sequenced by Sanger sequencing, and chromosomes were aligned to each other in Benchling. Table S2 shows the TLH and gene drive inheritance rate for each F1 parent. Average TLH was 3.6%, lower than the average 5.1% TLH seen in the F1 pools due to the necessity of overestimating SNP presence in the G3 sequence used for comparison to the pooled samples, but well within the F1 pool range (2.1-9.4% TLH in *zpg*-7280/N’Gousso F1 hybrids).

**Table S2:**
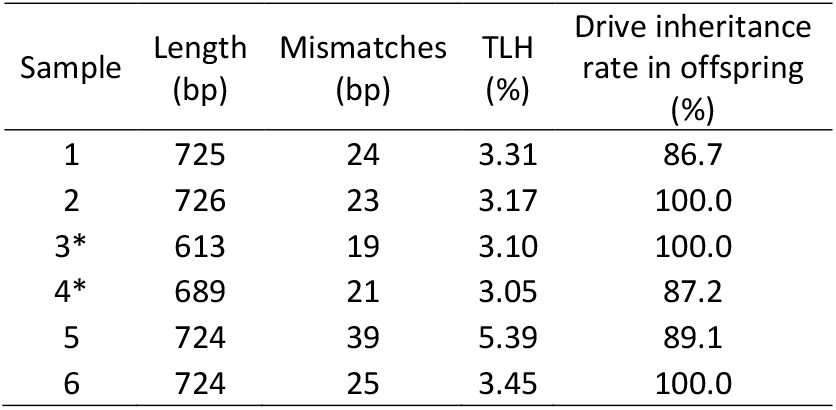
Target locus heterology (TLH) between chromosomes of zpg-7280/N’Gousso F1 hybrids, and the inheritance rate of the gene drive in their offspring, indicating the efficacy of the gene drive into a heterogeneous target chromosome. * sequences were truncated to omit poor quality sequences.

| Sample | Length (bp) | Mismatches (bp) | TLH (%) | Drive inheritance rate in offspring (%) |
| --- | --- | --- | --- | --- |
| 1 | 725 | 24 | 3.31 | 86.7 |
| 2 | 726 | 23 | 3.17 | 100.0 |
| 3* | 613 | 19 | 3.10 | 100.0 |
| 4* | 689 | 21 | 3.05 | 87.2 |
| 5 | 724 | 39 | 5.39 | 89.1 |
| 6 | 724 | 25 | 3.45 | 100.0 |

**Table S3:**
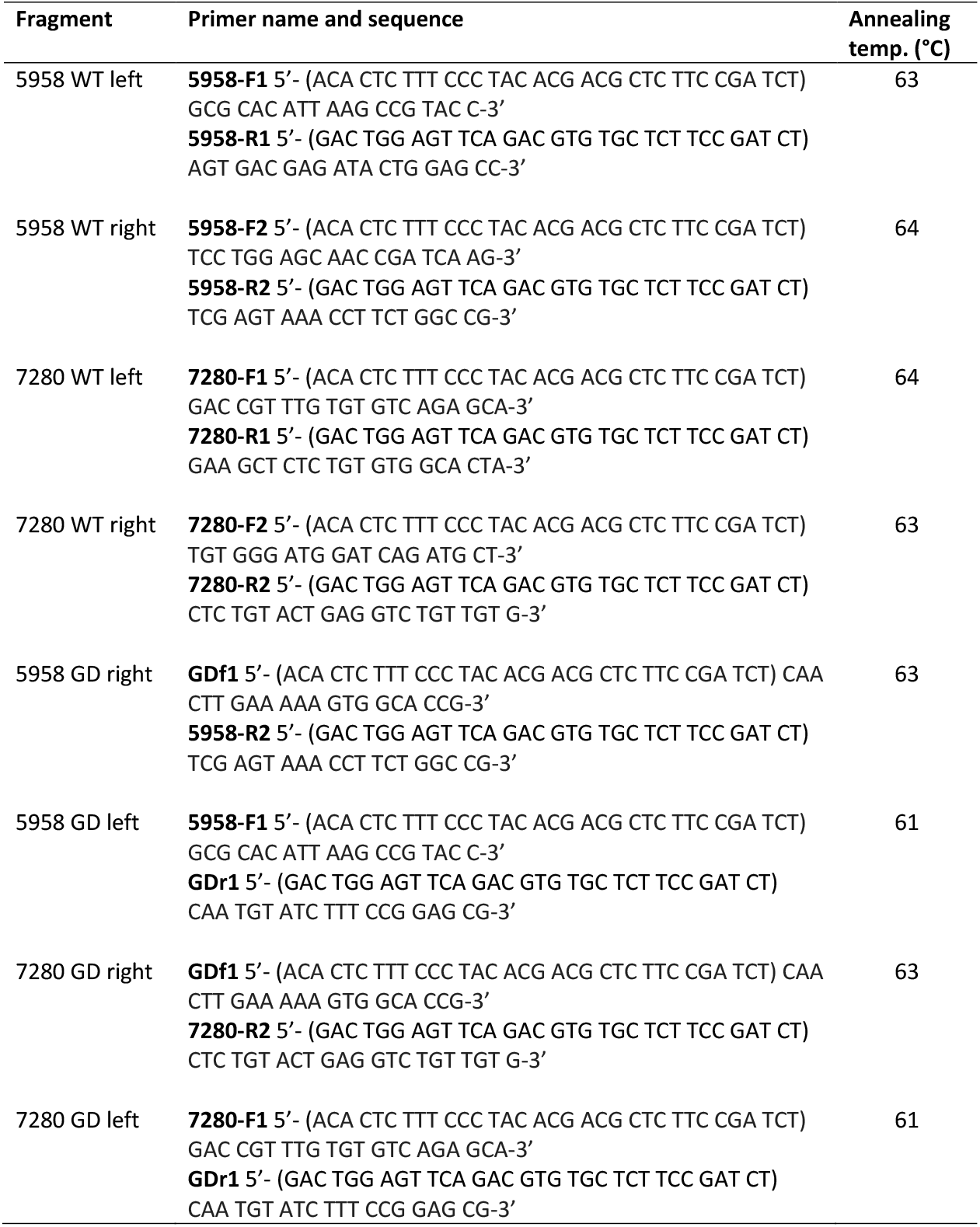
Primers used for amplicon sequencing of TLH regions. Sequences in brackets indicate Illumina adapter sequences (not included in annealing temperature calculation). WT = wild type, GD = gene drive.

**Figure S2:**
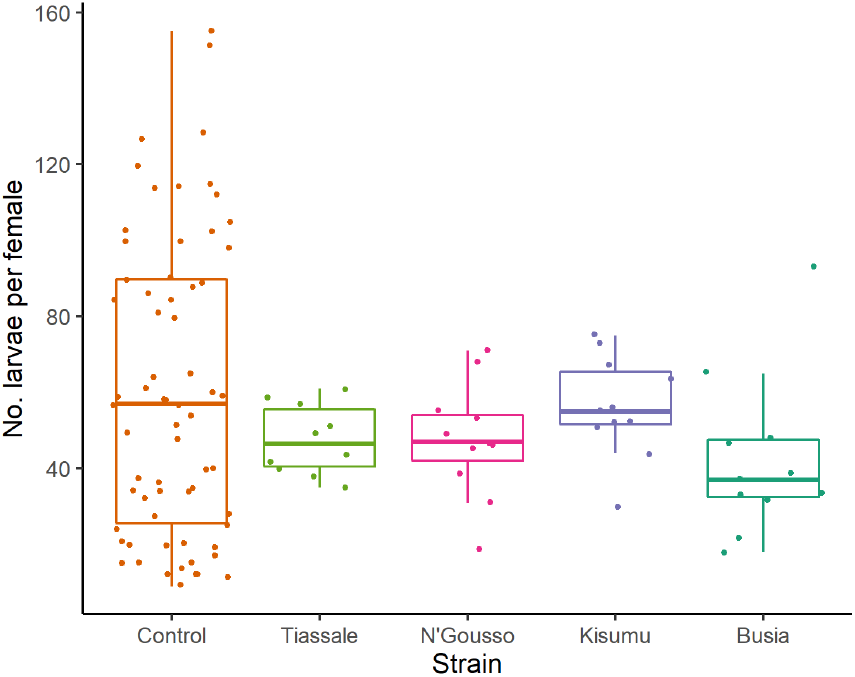
Comparison of larvae number produced by single females from hybrids of *zpg*-7280 and four different strains. Control larvae numbers were from Hammond *et al*. (2021) (n=66). Tiassale n=10, N’Gousso n=11, Kisumu n=11, Busia n=11. No significant difference in larval production was found between any strain (pairwise t test, p > 0.05 for all comparisons), suggesting that there was no reduction in fertility in gene drive/alternate strain hybrids. Hammond *et al*. (2021). Regulating the expression of gene drives is key to increasing their invasive potential and the mitigation of resistance. *PLoS Genetics* **17**(1), e1009321.

## References

1. World Health Organization. World Malaria Report 2022. Geneva; 2022.

2. Rodriguez MH. Residual Malaria: Limitations of Current Vector Control Strategies to Eliminate Transmission in Residual Foci. The Journal of Infectious Diseases 2021;223(Supplement_2):S55–S60, doi:10.1093/infdis/jiaa582

3. World Health Organization. Evaluation of genetically modified mosquitoes for the control of vector-borne diseases. Geneva; 2020.

4. World Health Organization.World malaria report 2020: 20 years of global progress and challenges. Geneva; 2020.

5. Burt A. Site-specific selfish genes as tools for the control and genetic engineering of natural populations. Proceedings of the Royal Society of London Series B: Biological Sciences 2003;270(1518):921–928

6. Gantz VM, Jasinskiene N, Tatarenkova O, et al. Highly efficient Cas9-mediated gene drive for population modification of the malaria vector mosquito Anopheles stephensi. Proceedings of the National Academy of Sciences 2015;112(49):E6736–E6743

7. Hammond A, Galizi R, Kyrou K, et al. A CRISPR-Cas9 gene drive system targeting female reproduction in the malaria mosquito vector Anopheles gambiae. Nature Biotechnology 2016;34(1):78–83, doi:10.1038/nbt.3439

8. Kyrou K, Hammond AM, Galizi R, et al. A CRISPR–Cas9 gene drive targeting doublesex causes complete population suppression in caged Anopheles gambiae mosquitoes. Nature Biotechnology 2018;36(11):1062–1066, doi:10.1038/nbt.4245

9. Windbichler N, Menichelli M, Papathanos PA, et al. A synthetic homing endonuclease-based gene drive system in the human malaria mosquito. Nature 2011;473(7346):212–215

10. Hoermann A, Habtewold T, Selvaraj P, et al. Gene drive mosquitoes can aid malaria elimination by retarding Plasmodium sporogonic development. Science Advances 2022;8(38):eabo1733, doi:doi:10.1126/sciadv.abo1733

11. Legros M, Marshall JM, Macfadyen S, et al. Gene drive strategies of pest control in agricultural systems: Challenges and opportunities. Evolutionary Applications 2021;14(9):2162–2178, doi:https://doi.org/10.1111/eva.13285

12. Scott MJ, Gould F, Lorenzen M, et al. Agricultural production: assessment of the potential use of Cas9-mediated gene drive systems for agricultural pest control. Journal of Responsible Innovation 2018;5(sup1):S98–S120, doi:10.1080/23299460.2017.1410343

13. Ceccaldi R, Rondinelli B, D’Andrea AD. Repair Pathway Choices and Consequences at the Double-Strand Break. Trends in Cell Biology 2016;26(1):52–64, doi:https://doi.org/10.1016/j.tcb.2015.07.009

14. Xue C, Greene EC. DNA Repair Pathway Choices in CRISPR-Cas9-Mediated Genome Editing. Trends in Genetics 2021;37(7):639–656, doi:https://doi.org/10.1016/j.tig.2021.02.008

15. Hammond AM, Kyrou K, Bruttini M, et al. The creation and selection of mutations resistant to a gene drive over multiple generations in the malaria mosquito. PLOS Genetics 2017;13(10):e1007039, doi:10.1371/journal.pgen.1007039

16. Beaghton AK, Hammond A, Nolan T, et al. Gene drive for population genetic control: non-functional resistance and parental effects. Proceedings of the Royal Society B: Biological Sciences 2019;286(1914):20191586, doi:doi:10.1098/rspb.2019.1586

17. Chang HH, Pannunzio NR, Adachi N, et al. Non-homologous DNA end joining and alternative pathways to double-strand break repair. Nature Reviews Molecular Cell Biology 2017;18(8):495–506

18. Unckless RL, Clark AG, Messer PW. Evolution of Resistance Against CRISPR/Cas9 Gene Drive. Genetics 2017;205(2):827–841, doi:10.1534/genetics.116.197285

19. Champer J, Reeves R, Oh SY, et al. Novel CRISPR/Cas9 gene drive constructs reveal insights into mechanisms of resistance allele formation and drive efficiency in genetically diverse populations. PLoS Genetics 2017;13(7):e1006796, doi:10.1371/journal.pgen.1006796

20. Champer J, Liu J, Oh SY, et al. Reducing resistance allele formation in CRISPR gene drive. Proceedings of the National Academy of Sciences 2018;115(21):5522–5527, doi:doi:10.1073/pnas.1720354115

21. Hammond A, Karlsson X, Morianou I, et al. Regulating the expression of gene drives is key to increasing their invasive potential and the mitigation of resistance. PLOS Genetics 2021;17(1):e1009321, doi:10.1371/journal.pgen.1009321

22. Marshall JM, Buchman A, Sánchez C HM, et al. Overcoming evolved resistance to population-suppressing homing-based gene drives. Scientific Reports 2017;7(1):3776, doi:10.1038/s41598-017-02744-7

23. Champer J, Yang E, Lee E, et al. A CRISPR homing gene drive targeting a haplolethal gene removes resistance alleles and successfully spreads through a cage population. Proceedings of the National Academy of Sciences 2020;117(39):24377–24383, doi:doi:10.1073/pnas.2004373117

24. Khatri BS, Burt A. A theory of resistance to multiplexed gene drive demonstrates the significant role of weakly deleterious natural genetic variation. Proceedings of the National Academy of Sciences 2022;119(32):e2200567119, doi:doi:10.1073/pnas.2200567119

25. Denholm I, Pickett JA, Devonshire AL, et al. Predicting insecticide resistance: mutagenesis, selection and response. Philosophical Transactions of the Royal Society of London Series B: Biological Sciences 1998;353(1376):1729–1734, doi:doi:10.1098/rstb.1998.0325

26. Doench JG, Fusi N, Sullender M, et al. Optimized sgRNA design to maximize activity and minimize off-target effects of CRISPR-Cas9. Nat Biotechnol 2016;34(2):184–191, doi:10.1038/nbt.3437

27. Elliott B, Jasin M. Repair of Double-Strand Breaks by Homologous Recombination in Mismatch Repair-Defective Mammalian Cells. Molecular and Cellular Biology 2001;21(8):2671–2682, doi:doi:10.1128/MCB.21.8.2671-2682.2001

28. Do AT, Brooks JT, Le Neveu MK, et al. Double-Strand Break Repair Assays Determine Pathway Choice and Structure of Gene Conversion Events in Drosophila melanogaster. G3 Genes|Genomes|Genetics 2014;4(3):425–432, doi:10.1534/g3.113.010074

29. Ang JXD, Nevard K, Ireland R, et al. Considerations for homology-based DNA repair in mosquitoes: Impact of sequence heterology and donor template source. PLoS Genetics 2022;18(2):e1010060

30. Clarkson CS, Miles A, Harding NJ, et al. Genome variation and population structure among 1142 mosquitoes of the African malaria vector species Anopheles gambiae and Anopheles coluzzii. Genome Research 2020;30(10):1533–1546

31. Coluzzi M, Sabatini A, Petrarca V, et al. Chromosomal differentiation and adaptation to human environments in the Anopheles gambiae complex. Transactions of the Royal Society of tropical Medicine and Hygiene 1979;73(5):483–497

32. Coluzzi M, Sabatini A, della Torre A, et al. A polytene chromosome analysis of the Anopheles gambiae species complex. Science 2002;298(5597):1415–1418

33. Gray EM, Rocca KA, Costantini C, et al. Inversion 2La is associated with enhanced desiccation resistance in Anopheles gambiae. Malaria Journal 2009;8(1):1–12

34. Manoukis NC, Powell JR, Touré MB, et al. A test of the chromosomal theory of ecotypic speciation in Anopheles gambiae. Proceedings of the National Academy of Sciences 2008;105(8):2940–2945

35. Petrarca V, Beier JC. Intraspecific chromosomal polymorphism in the Anopheles gambiae complex as a factor affecting malaria transmission in the Kisumu area of Kenya. The American Journal of Tropical Medicine and Hygiene 1992;46(2):229–237

36. Brooke B, Hunt R, Chandre F, et al. Stable chromosomal inversion polymorphisms and insecticide resistance in the malaria vector mosquito Anopheles gambiae (Diptera: Culicidae). Journal of Medical Entomology 2002;39(4):568–573

37. Ayala D, Zhang S, Chateau M, et al. Association mapping desiccation resistance within chromosomal inversions in the African malaria vector Anopheles gambiae. Molecular Ecology 2019;28(6):1333–1342

38. White BJ, Santolamazza F, Kamau L, et al. Molecular karyotyping of the 2La inversion in Anopheles gambiae. The American Journal of Tropical Medicine and Hygiene 2007;76(2):334–339

39. Powell J, Petrarca V, Della Torre A, et al. Population structure, speciation, and introgression in the Anopheles gambiae complex. Parassitologia 1999;41(1-3):101–113

40. Sharakhov IV, White BJ, Sharakhova MV, et al. Breakpoint structure reveals the unique origin of an interspecific chromosomal inversion (2La) in the Anopheles gambiae complex. Proceedings of the National Academy of Sciences 2006;103(16):6258–6262

41. Fouet C, Gray E, Besansky NJ, et al. Adaptation to aridity in the malaria mosquito Anopheles gambiae: chromosomal inversion polymorphism and body size influence resistance to desiccation. PloS ONE 2012;7(4):e34841

42. Kirkpatrick M. How and why chromosome inversions evolve. PLoS Biology 2010;8(9):e1000501

43. Crown KN, Miller DE, Sekelsky J, et al. Local Inversion Heterozygosity Alters Recombination throughout the Genome. Current Biology 2018;28(18):2984–2990.e3, doi:https://doi.org/10.1016/j.cub.2018.07.004

44. Lucchesi JC, Suzuki DT. The interchromosomal control of recombination. Annual Review of Genetics 1968;2(1):53–86, doi:10.1146/annurev.ge.02.120168.000413

45. Hunter CM, Huang W, Mackay TFC, et al. The Genetic Architecture of Natural Variation in Recombination Rate in Drosophila melanogaster. PLOS Genetics 2016;12(4):e1005951, doi:10.1371/journal.pgen.1005951

46. Stapley J, Feulner PGD, Johnston SE, et al. Variation in recombination frequency and distribution across eukaryotes: patterns and processes. Phil Trans R Soc B 2017;372(1736):20160455, doi:https://doi.org/10.1098/rstb.2016.0455

47. Guichard A, Haque T, Bobik M, et al. Efficient allelic-drive in Drosophila. Nature Communications 2019;10(1):1640, doi:10.1038/s41467-019-09694-w

48. Stump A, Pombi M, Goeddel L, et al. Genetic exchange in 2La inversion heterokaryotypes of Anopheles gambiae. Insect Molecular Biology 2007;16(6):703–709

49. Hoermann A, Tapanelli S, Capriotti P, et al. Converting endogenous genes of the malaria mosquito into simple non-autonomous gene drives for population replacement. eLife 2021;10(e58791, doi:10.7554/eLife.58791

50. Ellis DA, Avraam G, Hoermann A, et al. Testing non-autonomous antimalarial gene drive effectors using self-eliminating drivers in the African mosquito vector Anopheles gambiae. PLOS Genetics 2022;18(6):e1010244, doi:10.1371/journal.pgen.1010244

51. Champer J, Wen Z, Luthra A, et al. CRISPR Gene Drive Efficiency and Resistance Rate Is Highly Heritable with No Common Genetic Loci of Large Effect. Genetics 2019;212(1):333–341, doi:10.1534/genetics.119.302037

52. Courret C, Chang C-H, Wei KH-C, et al. Meiotic drive mechanisms: lessons from Drosophila. Proc R Soc B 2019;286(1913), doi:https://doi.org/10.1098/rspb.2019.1430

53. Claycomb JM, Benasutti M, Bosco G, et al. Gene Amplification as a Developmental Strategy: Isolation of Two Developmental Amplicons in Drosophila. Developmental Cell 2004;6(1):145–155, doi:https://doi.org/10.1016/S1534-5807(03)00398-8

54. Lee Y, Collier TC, Sanford MR, et al. Chromosome Inversions, Genomic Differentiation and Speciation in the African Malaria Mosquito Anopheles gambiae. PLoS ONE 2013;8(3):1–12, doi:10.1371/journal.pone.0057887

55. Hong CC, Hashimoto C. The maternal nudel protein of Drosophila has two distinct roles important for embryogenesis. Genetics 1996;143(4):1653–61, doi:10.1093/genetics/143.4.1653

56. Bateman JR, Lee AM, Wu C-T. Site-specific transformation of Drosophila via ϕC31 integrase-mediated cassette exchange. Genetics 2006;173(2):769–777

57. Harris C, Lambrechts L, Rousset F, et al. Polymorphisms in Anopheles gambiae Immune Genes Associated with Natural Resistance to Plasmodium falciparum. PLOS Pathogens 2010;6(9):e1001112, doi:10.1371/journal.ppat.1001112

58. Edi CV, Koudou BG, Jones CM, et al. Multiple-insecticide resistance in Anopheles gambiae mosquitoes, Southern Côte d’Ivoire. Emerg Infect Dis 2012;18(9):1508–11, doi:10.3201/eid1809.120262

59. Reid J. Pupal differences between species A and B of the Anopheles gambiae group from Kisumu, East Africa. Mosquito Systematics 1975;

60. Clement K, Rees H, Canver MC, et al. CRISPResso2 provides accudate and rapid genome editing sequence analysis. Nature Biotechnology 2019;37(3):224–226, doi:10.1038/s41587-019-0032-3

61. Reid W, Williams AE, Sanchez-Vargas I, et al. Assessing single-locus CRISPR/Cas9-based gene drive variants in the mosquito Aedes aegypti via single-generation crosses and modeling. G3 Genes|Genomes|Genetics 2022;12(12), doi:10.1093/g3journal/jkac280

62. Holt RA, Subramanian GM, Halpern A, et al. The Genome Sequence of the Malaria Mosquito Anopheles gambiae. Science 2002;298(5591):129–149, doi:doi:10.1126/science.1076181

63. Clarkson CS, Weetman D, Essandoh J, et al. Adaptive introgression between Anopheles sibling species eliminates a major genomic island but not reproductive isolation. Nature Communications 2014;5(1):4248, doi:10.1038/ncomms5248

64. Pollegioni P, Persampieri T, Minuz RL, et al. Introgression of a synthetic sex ratio distortion transgene into different genetic backgrounds of Anopheles coluzzii. Insect Molecular Biology 2023;32(1):56–68, doi:https://doi.org/10.1111/imb.12813

65. The Anopheles gambiae 1000 Genomes Consortium. Ag1000G. https://www.malariagen.net/mosquito/ag1000g; 2014.

66. Noble C, Min J, Olejarz J, et al. Daisy-chain gene drives for the alteration of local populations. Proceedings of the National Academy of Sciences 2019;116(17):8275–8282, doi:doi:10.1073/pnas.1716358116

67. Dhole S, Lloyd AL, Gould F. Tethered homing gene drives: A new design for spatially restricted population replacement and suppression. Evolutionary Applications 2019;12(8):1688–1702, doi:https://doi.org/10.1111/eva.12827

68. Willis K, Burt A. Double drives and private alleles for localised population genetic control. PLOS Genetics 2021;17(3):e1009333, doi:10.1371/journal.pgen.1009333

69. Garrood WT, Kranjc N, Petri K, et al. Analysis of off-target effects in CRISPR-based gene drives in the human malaria mosquito. Proceedings of the National Academy of Sciences 2021;118(22):e2004838117

70. James SL, Marshall JM, Christophides GK, et al. Toward the definition of efficacy and safety criteria for advancing gene drive-modified mosquitoes to field testing. Vector-Borne and Zoonotic Diseases 2020;20(4):237–251

71. Long KC, Alphey L, Annas GJ, et al. Core commitments for field trials of gene drive organisms. Science 2020;370(6523):1417–1419, doi:doi:10.1126/science.abd1908

72. Lynd et al. (2019). LLIN Evaluation in Uganda Project (LLINEUP): a cross-sectional survey of species diversity and insecticide resistance in 48 districts of Uganda. Parasites & Vectors 12(1), 1–10.

