## Supplementary material for "Measuring the impact of genetic heterogeneity and chromosomal inversions on the efficacy of CRISPR-Cas9 gene drives in different strains of *Anopheles gambiae*": Table S1

### Supporting information

**S1 Appendix: Analysis of a fourth strain, Busia**

In a separate piece of work, a fourth strain Busia (*Anopheles gambiae* s.s., captured in Uganda in 2018) (Lynd, *et al.,* 2019) was analysed for homing rates in F1 hybrids with both *vas2*-5958 and *zpg*-7280. Male hybrids of Busia with both gene drives were used in *en masse* crosses with wild type G3 females; females were put in single deposition for drive rate analysis (four from crosses with Busia/*vas2*-5958 hybrids, and 11 from crosses with Busia/*zpg*-7280 hybrids). Target site heterology (TLH) was calculated from wild type Busia using Sanger sequencing, with nine replicates on the left-hand side of the *vas2*-5958 cut site, and three replicates spanning 700 bp either side of the *zpg*-7280 cut site (accession: PRJNA914102). While the Busia samples were processed differently to the remaining three strains and were therefore left out of the main analysis, they show the same pattern of uninterrupted homing regardless of TLH, indicating the robustness of this effect to different analysis methods. All supporting information will include results from the Busia strain alongside Kisumu, N'Gousso and Tiassale.

Lynd *et al.* (2019). LLIN Evaluation in Uganda Project (LLINEUP): a cross-sectional survey of species diversity and insecticide resistance in 48 districts of Uganda. *Parasites & Vectors* **12**(1), 1-10.

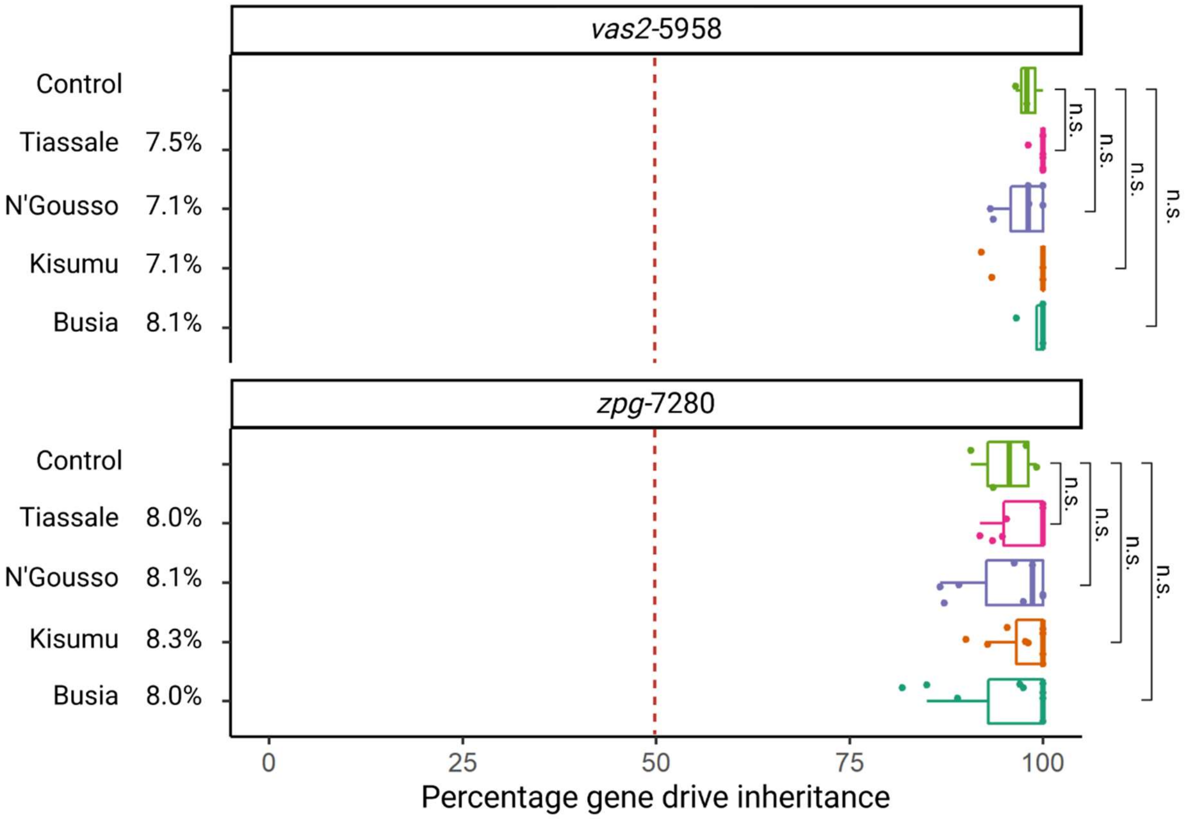


**Figure S1: Gene drive inheritance rate analysis of Kisumu, N'Gousso, Tiassale and Busia hybrids with *zpg*-7280 and *vas2*-5958 gene drives.** TLH is given as a percentage next to each strain name; statistical analysis was conducted using a pairwise Wilcoxon test with false discovery rate correction. n.s. – non-significant.

**Table S1: Pairwise Wilcoxon test p values with false discovery rate correction for gene drive inheritance rate.** Comparisons were performed for hybrids with all four strains of both gene drives. No comparisons were statistically significant.

| ***vas2*-5958** | | | | | ***zpg*-7280** | | | | |
|---|---|---|---|---|---|---|---|---|---|
| | *Busia* | Kisumu | N'Gousso | Tiassale | | *Busia* | Kisumu | N'Gousso | Tiassale |
| Kisumu | *1.00* | - | - | - | Kisumu | *0.91* | - | - | - |
| N'Gousso | *0.55* | 0.55 | - | - | N'Gousso | *1.00* | 0.91 | - | - |
| Tiassale | *0.55* | 0.55 | 0.17 | - | Tiassale | *0.91* | 1.00 | 0.91 | - |
| Control | *0.55* | 0.55 | 1.00 | 0.17 | Control | *0.91* | 0.91 | 0.91 | 0.91 |

**Table S2: Primers used for amplicon sequencing of TLH regions**. Sequences in brackets indicate Illumina adapter sequences (not included in annealing temperature calculation). WT = wild type, GD = gene drive.

| Fragment | Primer name and sequence | Annealing temp. (°C) |
|---|---|---|
| 5958 WT left | **5958-F1** 5'- (ACA CTC TTT CCC TAC ACG ACG CTC TTC CGA TCT) GCG CAC ATT AAG CCG TAC C-3'<br>**5958-R1** 5'- (GAC TGG AGT TCA GAC GTG TGC TCT TCC GAT CT) AGT GAC GAG ATA CTG GAG CC-3' | 63 |
| 5958 WT right | **5958-F2** 5'- (ACA CTC TTT CCC TAC ACG ACG CTC TTC CGA TCT) TCC TGG AGC AAC CGA TCA AG-3'<br>**5958-R2** 5'- (GAC TGG AGT TCA GAC GTG TGC TCT TCC GAT CT) TCG AGT AAA CCT TCT GGC CG-3' | 64 |
| 7280 WT left | **7280-F1** 5'- (ACA CTC TTT CCC TAC ACG ACG CTC TTC CGA TCT) GAC CGT TTG TGT GTC AGA GCA-3'<br>**7280-R1** 5'- (GAC TGG AGT TCA GAC GTG TGC TCT TCC GAT CT) GAA GCT CTC TGT GTG GCA CTA-3' | 64 |
| 7280 WT right | **7280-F2** 5'- (ACA CTC TTT CCC TAC ACG ACG CTC TTC CGA TCT) TGT GGG ATG GAT CAG ATG CT-3'<br>**7280-R2** 5'- (GAC TGG AGT TCA GAC GTG TGC TCT TCC GAT CT) CTC TGT ACT GAG GTC TGT TGT G-3' | 63 |
| 5958 GD right | **GDf1** 5'- (ACA CTC TTT CCC TAC ACG ACG CTC TTC CGA TCT) CAA CTT GAA AAA GTG GCA CCG-3'<br>**5958-R2** 5'- (GAC TGG AGT TCA GAC GTG TGC TCT TCC GAT CT) TCG AGT AAA CCT TCT GGC CG-3' | 63 |
| 5958 GD left | **5958-F1** 5'- (ACA CTC TTT CCC TAC ACG ACG CTC TTC CGA TCT) GCG CAC ATT AAG CCG TAC C-3'<br>**GDr1** 5'- (GAC TGG AGT TCA GAC GTG TGC TCT TCC GAT CT) CAA TGT ATC TTT CCG GAG CG-3' | 61 |
| 7280 GD right | **GDf1** 5'- (ACA CTC TTT CCC TAC ACG ACG CTC TTC CGA TCT) CAA CTT GAA AAA GTG GCA CCG-3'<br>**7280-R2** 5'- (GAC TGG AGT TCA GAC GTG TGC TCT TCC GAT CT) CTC TGT ACT GAG GTC TGT TGT G-3' | 63 |
| 7280 GD left | **7280-F1** 5'- (ACA CTC TTT CCC TAC ACG ACG CTC TTC CGA TCT) GAC CGT TTG TGT GTC AGA GCA-3'<br>**GDr1** 5'- (GAC TGG AGT TCA GAC GTG TGC TCT TCC GAT CT) CAA TGT ATC TTT CCG GAG CG-3' | 61 |

**Table S3: TLH for Kisumu, N'Gousso, Tiassale and Busia compared to G3.** TLH was calculated by comparing combined sequences from each strain to wild type G3 sequences and calculating the percentage of base pairs differing from the wild type. For the additional fourth strain, Busia, only the 309 bp sequence to the left of the *vas2*-5958 cut site was used to calculate TLH.

| ***vas2*-5958 target site heterology** | | | | | |
|---|---|---|---|---|---|
| | **# in pool** | **# polymorphic sites** | **# bp different** | **Total bp of site** | **% of bp different** |
| Kisumu | 37 | 39 | 45 | 637 | 7.1 |
| N'Gousso | 33 | 38 | 45 | 637 | 7.1 |
| Tiassale | 34 | 42 | 48 | 637 | 7.5 |
| *Busia ** | *9* | *19* | *25* | *309* | *8.1* |
| ***zpg*-7280 target site heterology** | | | | | |
| | **# in pool** | **# polymorphic sites** | **# bp different** | **Total bp of site** | **% of bp different** |
| Kisumu | 37 | 51 | 57 | 690 | 8.3 |
| N'Gousso | 33 | 50 | 56 | 690 | 8.1 |
| Tiassale | 34 | 49 | 55 | 690 | 8.0 |
| *Busia* | *3* | *55* | *55* | *690* | *8.0* |

**Figure S2: Comparison of larvae number produced by single females from hybrids of *zpg*-7280 and four different strains**. Control larvae numbers were from Hammond *et al.* (2021) (n=66). Tiassale n=10, N'Gousso n=11, Kisumu n=11, Busia n=11. No significant difference in larval production was found between any strain (pairwise t test, p > 0.05 for all comparisons), suggesting that there was no reduction in fertility in gene drive/alternate strain hybrids.

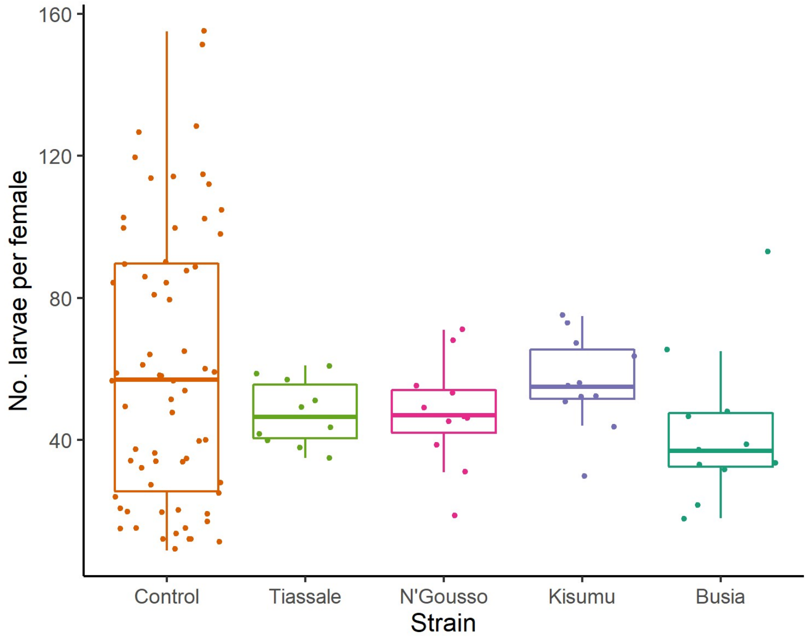



Hammond *et al.* (2021). Regulating the expression of gene drives is key to increasing their invasive potential and the mitigation of resistance. *PLoS Genetics* **17**(1), e1009321.
